# Synaptic editing of frontostriatal circuitry prevents excessive grooming in SAPAP3-deficient mice

**DOI:** 10.1101/2025.03.27.645613

**Authors:** Kathryn K. Walder-Christensen, Hannah A. Soliman, Nicole Calakos, Kafui Dzirasa

## Abstract

Synaptic dysfunction has been implicated as a key mechanism underlying the pathophysiology of psychiatric disorders. Most pharmacological therapeutics for schizophrenia, autism spectrum disorder, obsessive-compulsive disorder, and major depressive disorder temporarily augment chemical synapse function. Nevertheless, medication non-compliance is a major clinical challenge, and behavioral dysfunction often returns following pharmacotherapeutic discontinuation. Here, we deployed a designer electrical synapse to edit a single class of chemical synapses in a genetic mouse model of obsessive-compulsive disorder (OCD). Editing these synapses in juvenile mice normalized circuit function and prevented the emergence of pathological repetitive behavior in adulthood. Thus, we establish precision circuit editing as a putative strategy for preventative psychotherapeutics.

---

Alterations to synapse function have been identified across many neuropsychiatric disorders^1-3^. Multiple monogenic mutations that cause synaptic dysfunction have been identified in humans with psychiatric disorders^4^. In other cases, rare monogenic mutations causally implicated in synaptic dysfunction have been identified across psychiatric disorders with shared behavioral phenotypes. For both instances, myriad studies have modeled these monogenic mutations in murine organisms^5,6^. One such example is the *Sapap3*-/- mouse model.

Sapap3 is a postsynaptic scaffolding protein that is highly expressed in the cortex and striatum^7,8^. The protein is localized to the PSD of excitatory synapses and binds PSD95 and SHANK proteins to coordinate ion channels to the underlying cytoskeleton^9^. Due to this key structural role in excitatory synapses, deficits in *Sapap3* yield altered synaptic structure and function^7^. Although the gene does not reach genome wide significance in human disorders, *Sapap3* dysfunction has been associated with neuropsychiatric disorders characterized by repetitive behavior including OCD, trichotillomania, Tourette syndrome, and autism spectrum disorder^9^.

A plethora of studies have investigated brain connectivity differences in patients with OCD, indicating several regions of interest. Neuroimaging studies point to anatomical and functional differences in orbitofrontal cortex (OFC) and/or striatum (Str) in patients with OCD ^10-12^. Indeed, functional connectivity studies in patients with OCD have revealed altered connectivity of OFC in subjects with OCD and their first-degree relatives, pointing to an underlying genetically mediated process^13^. Further studies have revealed altered functional connectivity between orbitofrontal-striatal regions ^14-16^, indicating the contributions of the OFC and Str towards behavioral disruption.

OCD is characterized by repetitive behaviors such as excessive hand washing. In some cases, this hand washing can be so severe that it yields skin lesions. Similarly, mice lacking *Sapap3* (*Sapap3* -/- mice) exhibit excessive facial grooming, yielding severe facial lesions compared to healthy littermate controls^7^. OFC and Str dysfunction have been implicated in this behavioral phenotype displayed by *Sapap3*-/- mice. *Sapap3*-/- mice exhibit increased inhibitory neuronal activity in the OFC, and optogenetic stimulation of neuronal activity in OFC or of OFC terminals within Str suppresses their pathological grooming^17^. Within striatum, virally mediated expression of Sapap3 during early development reduces grooming bouts, lesion severity, and anxiety-related behavior in adult *Sapap3*-/- mice^7^, and optogenetic stimulation of Str parvalbumin-expressing (PV+) inhibitory neurons acutely suppresses their pathological grooming^18^. This latter stimulus response is enhanced if light stimulation is timed to LFP features in OFC that signal grooming onset^18^.

While together, these studies implicate a circuit composed of OFC excitatory projection neurons and Str PV+ interneurons in the grooming phenotype displayed by *Sapap3*-/- mice, definitive evidence for such a pathophysiological mechanism remains lacking. Excitatory projection neurons from OFC form chemical synapses with striatal projection neurons (SPNs, either expressing D1 or D2 receptors) and PV+ interneurons of the dorsal striatum (Figure 1C; Supplemental Fig. S1)^19^. Thus, stimulating OFC terminals in Str drives all three of these subcircuits (OFC⟶Str D1+ SPN, OFC⟶Str D2+ SPN, and OFC⟶Str PV). Timing the stimulation of PV+ interneurons to OFC LFP features that signal grooming onset enhances the potency of this manipulation^18^, but it has yet to be established that OFC is the only brain structure that can optimize the behavioral effect of Str PV+ interneuron stimulation. This is a particular concern since OFC LFP activity shows a high degree of spectral synchrony with other brain regions^20,21^, and other regions also project to striatum and form synapses with Str PV+ interneurons. Thus, it could be argued that other circuits may also be central to the induction of pathological grooming. Indeed, to directly test this question as to whether deficits in the OFC⟶Str PV+ circuit specifically mediate pathological grooming in the *Sapap3*-/- mice, one must selectively target this circuit while leaving the OFC⟶Str D1+ SPN and OFC⟶Str D2+ SPN circuits unaltered. Yet, tools that selectively modify the interaction between a pair of cell types remain scarce.

**Figure 1.**
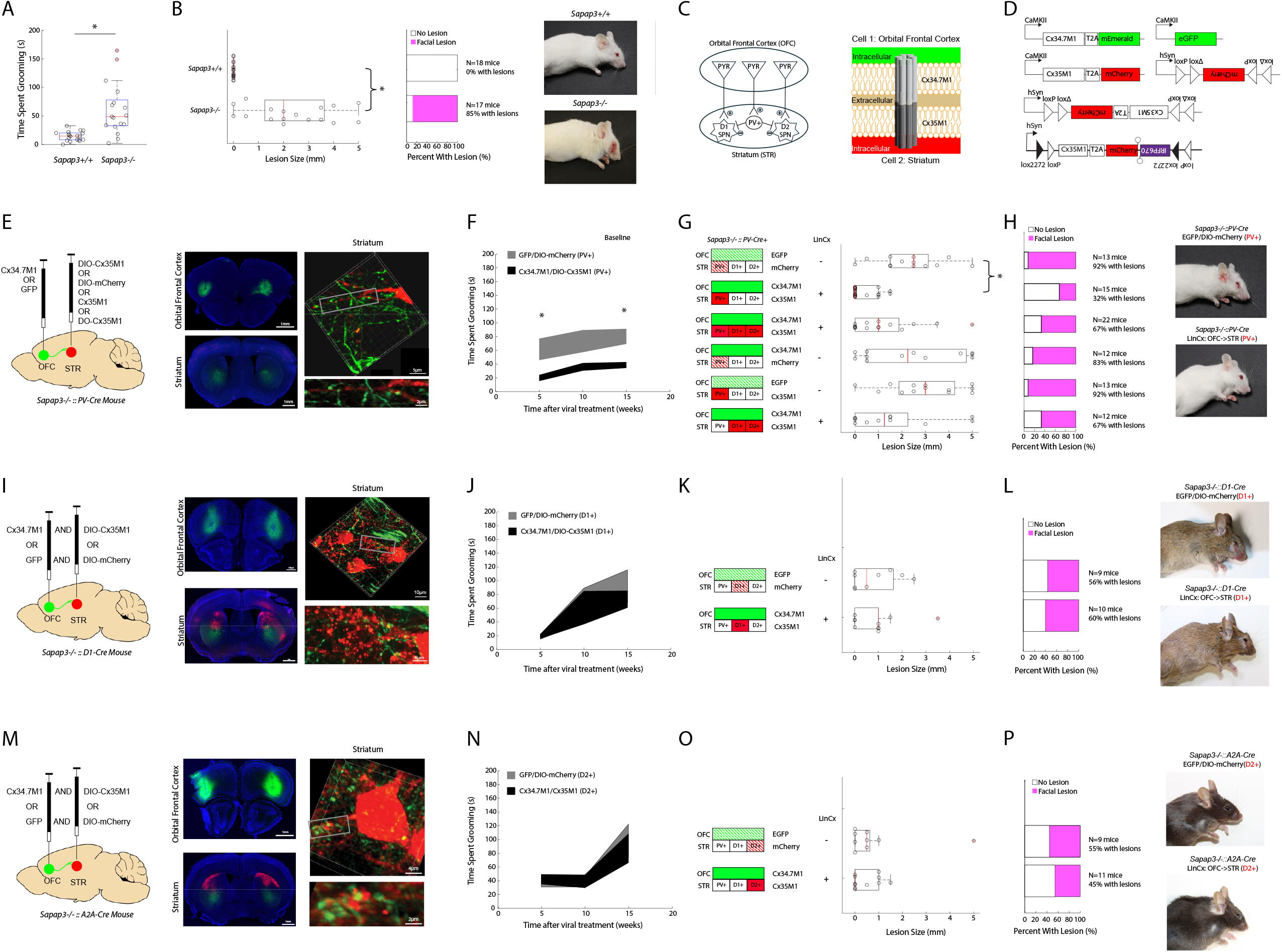
LinCx expression in the OFC⟶Str-PV interneuron circuit selectively reduces pathological grooming in *Sapap3*-/- mice. **A)** Quantification of grooming in *Sapap3*-/- mice and their littermate controls (*p<0.001 using Mann-Whitney test). **B)** Facial lesion size in *Sapap3*-/- and littermate controls at age 22 weeks (left, *p<0.001 using Mann-Whitney test). The percentage of mice with facial lesions is shown in the center, and representative images of *Sapap3*+/+ and *Sapap3*-/- mice are shown to the right. **C)** Schematic of targeted OFC⟶Str circuitry involving excitatory pyramidal neuron synapses onto each of 3 main cell types in Str: D1+ SPNs, D2+ SPNs, and PV+ interneurons. Within Str, PV+ interneurons synapse onto D1 and D2 SPNs and each other (left). Schematic of intended heteromeric gap junction. Green represents mEmerald/EGFP, while red represents mCherry. The Cx34.7_M1_ (light grey) hemichannel and mEmerald are expressed in OFC projection neurons. Striatal neurons express the reciprocal hemichannel, Cx35_M1_ (dark grey) and mCherry. Together, the Cx34.7_M1_ and Cx35_M1_ constitute a heterotypic gap junction, creating a connection between the intracellular space of OFC projection neurons and striatal neurons (right). **D)** Diagram of AAV9 viral constructs used to target OFC and Str. **E)** Diagram of viral manipulation strategy into OFC and Str in PV-Cre mice. AAV-Cx34.7_M1_ or AAV9-GFP was injected into OFC bilaterally. Additionally, AAV-DIO-Cx35_M1_, AAV-DO-Cx35_M1_, or AAV-DIO-mCherry was injected into Str bilaterally (left). Representative image of LinCx viral expression. Green fluorescence indicates AAV-CamK2-Cx34.7_M1_-T2A-mEmerald expression in OFC soma, and in OFC terminals in Str. Red fluorescence indicates Str soma that express AAV-hSyn-DIO-Cx35_M1_-T2A-mCherry (center). A confocal rendering of OFC (green) projections contacting PV+ interneurons (red) in Str in a *Sapap3-/- ::PV-Cre* mouse is also shown (right-grey box enlarged on bottom right). **F)** Quantification of grooming in *Sapap3-/- ::PV-Cre* mice following OFC-STR PV+ circuit editing (*p<0.05 using mixed effects model ANOVA followed by Dunn’s correction; note that other control groups are shown in Supplemental Fig. S2). Data shown as mean ± s.e.m. **G)** Quantification of facial lesions in *Sapap3-/- ::PV-Cre* mice 15 weeks post-viral manipulation (*p<0.0001 using Kruskal-Wallis test). The middle “LinCx” column indicates which rows are tests of LinCx manipulations (+) vs. hemichannel or fluorophore-only control groups (-). **H)** The percentage of *Sapap3-/- ::PV-Cre* mice with facial lesions is shown for each group (left), and representative images of a LinCx-edited *Sapap3-/- ::PV-Cre* mouse and a non-edited control mouse are shown (right). **I)** Viral injection strategy into OFC and striatum of *Sapap3-/- ::D1-Cre* mice. Representative image of LinCx viral expression (center) and confocal rendering of OFC (green) projections contacting D1-expressing striatal projection neurons (red) in Str in a *Sapap3*-/- ::D1-Cre mouse (right-grey box enlarged on bottom right). **J)** Quantification of grooming p=0.9098 for viral group x time effect, p=0.8635 for viral group effect, and p=0.0012 for time effect using mixed effects model; data shown as mean ± s.e.m.) and **K)** facial lesions in viral-manipulated mice (p=0.95 using Mann-Whitney test). **L)** The percentage of *Sapap3-/- ::D1-Cre* mice with facial lesions is shown for each group (left), and representative images of a LinCx-edited mouse and a non-edited control mouse are shown (right). **M)** Viral injection strategy into OFC and Str of *Sapap3-/- ::A2A-Cre* mice. Representative image of LinCx viral expression (center) and confocal rendering of OFC (green) projections contacting A2A-expressing SPNs (red) in Str in a *Sapap3-/- ::A2A-Cre* mouse (right-grey box enlarged on bottom right). **N)** Quantification of grooming (P=0.8765 for viral group x time effect using mixed effects model; data shown as mean ± s.e.m.) and **O)** facial lesions in viral-manipulated mice (p=0.95 using Mann Whitney test). **P)** The percentage of *Sapap3-/- ::A2A-Cre* mice with facial lesions is shown for each group (left), and representative images of a LinCx-edited mouse and a non-edited control mouse are shown (right). For box and whisker plots, the central mark is the median, the edges of the box are the 25th and 75th percentiles, the whiskers extend to the most extreme datapoints the algorithm considers to be not outliers, and the outliers are plotted individually as a “+”.

We have recently created a tool called <u>L</u>ong-term <u>In</u>tegration of circuits using <u>C</u>onne<u>x</u>ins (LinCx). LinCx consists of two proteins, Cx34.7_M1_ and Cx35_M1_, which are designed to exclusively dock with each other and in doing so, form a functional gap junction^22^. Each of the proteins can be targeted to distinct cell membranes that are in close proximity to create an electrical synapse between the pair of cells. Thus, LinCx augments the function of chemical synapses by creating an adjacent electrical synapse. Here, we first targeted LinCx to the circuit involving OFC projection neurons and striatal PV+ interneurons. We crossed PV-Cre mice with *Sapap3*-/-, to generate *Sapap3-/- ::PV-Cre* mice. We then generated connexin-T2A-fluorophore expressing viruses such that an epitope-tagged connexin and soluble fluorophore would be produced (Fig. 1D). The Cx34.7_M1_ construct was designed under the CaMKIIα promoter (AAV-CaMKII-Cx34.7_M1_-2xHA-T2A-mEmerald), such that it would be expressed in neurons that project to Str and most other neuron types. The Cx35 construct was doubly floxed such that it would only express protein in the presence of Cre recombinase (AAV-hSyn-DIO-Cx35_M1_-FLAG-T2A-mCherry).

We first verified that adult *Sapap3*-/- mice exhibit excessive grooming and facial lesions (Fig. 1A-B, U=36, p<0.001 for comparison of grooming time and U=18, p<0.001 for comparison of lesions size between 22-week-old *Sapap3*+/+ and *Sapap3*-/- mice using a two-tailed Mann-Whitney U test; n=18/17). We then injected AAV-CaMKII-Cx34.7_M1_-2xHA-T2A-mEmerald in OFC and the AAV-DIO-Cx35_M1_-FLAG-T2A-mCherry in Str of *Sapap3-/- ::PV-Cre* mice (Fig. 1C-D); we will refer to these mice as “LinCx-edited mice” moving forward. Control mice were injected with similar viral constructs except for the replacement of LinCx proteins with control fluorophores, AAV-CaMKII-EGFP in OFC and AAV-DIO-mCherry in Str. We also tested 4 additional control groups with other unique patterns of Cx34.7_M1_ or Cx35_M1_ expression in OFC and/or Str (described further below in the next paragraph). We were able to identify green processes (from Cx34.7_M1_ expressing cells in OFC) adjacent to red processes (Cx35_M1_ expressing cells in Str; see Fig. 1E), indicating the potential for each component of LinCx to be in close enough proximity to form functional gap junctions. *Sapap3-/- ::PV-Cre* mice injected with Cx34.7_M1_ in OFC and DIO-Cx35_M1_ in Str were assessed for grooming behavior at 5, 10, and 15 weeks after viral infection (F_1.868, 145.7_=30.88, p<0.0001 for virus group × time effect for comparison across all 6 viral groups using mixed-effects model ANOVA; Fig. 1F). These mice showed a significant decrease in grooming time at both 5 and 15 weeks, compared to control mice (Dunn’s multiple comparison test Z=2.845, adjusted p-value= 0.0222 at 5 weeks; Z=1.773, adjusted p-value=0.3809 at 10 weeks, and Z=2.760, adjusted p-value= 0.0289 at 15 weeks; Fig. 1F; see also Supplemental Fig. S2). Furthermore, this manipulation markedly decreased both the lesion size and the emergence of facial lesions (lesion size: H=28.21, p<0.0001 for comparison of lesion size across all 6 viral groups using Kruskal-Wallis test; post-hoc Dunn’s multiple comparison test between fluorophore control and LinCx-edited mice: Z=3.807, adjusted p-value = 0.0007; Fig. 1G-H, see row 1 and 2).

We then performed control experiments to verify that both halves of the gap junction were needed for this suppression of excessive grooming. Here, mice were injected with either CaMKII-Cx34.7_M1_-2xHA-T2A-mEmerald into OFC or DIO-Cx35_M1_-FLAG-T2A-mCherry into Str. In comparison to the dual fluorophore control, there was no significant difference in grooming time at 5, 10, or 15 weeks for either connexin independently (Dunn’s multiple comparison test: Z=0.6029, adjusted p-value=>0.9999 at 5 weeks; Z=0.3016, adjusted p-value=>0.9999 at 10 weeks; Z= 0.8154, adjusted p-value=>0.9999 at 15 weeks all for OFC-Cx34.7_M1_-2xHA-T2A-mEmerald. Dunn’s multiple comparison test Z=0.5668, adjusted p-value=>0.9999 at 5 weeks; Z=0.8961, adjusted p-value=>0.9999 at 10 weeks; Z=0.1408, adjusted p-value=>0.9999 at 15 weeks all for Str-DIO-Cx35_M1_; Supplemental Fig. S2A). Similarly, compared to the control mice, we observed no difference in lesion size when only one connexin was expressed (Dunn’s multiple comparison test Z=0.3660, adjusted p-value>0.9999 for OFC-CaMKII-Cx34.7_M1_; Z=0.6533, adjusted p-value >0.9999 for Str-DIO-Cx35_M1_; Fig. 1G-H, see row 4 and 5). We also tested whether selective expression of LinCx across multiple OFC-Str synaptic targets was required to suppress excessive grooming. We created a Cx35_M1_ virus that targeted multiple types of neurons (AAV-CaMKII-Cx35_M1_-FLAG-T2A-mCherry), and injected the Str of *Sapap3-/- ::PV-Cre* mice (see Supplemental Fig. S3). Mice were also injected with the CaMKII-Cx34.7_M1_ virus in OFC. With this strategy, we observed no statistical differences in grooming time at 5-15 weeks, compared to the control mice which expressed only fluorophores (Dunn’s multiple comparison test Z=2.335, adjusted p-value=0.0978 at 5 weeks; Dunn’s multiple comparison test Z=0.9569, adjusted p-value=>0.9999 at 15 Week all for AAV-CaMKII-Cx35_M1_; Supplemental Fig. S2B). Similarly, we found no significant decreases in their lesion size after 15 weeks of viral expression (Dunn’s multiple comparison test Z=1.925, adjusted p-value=0.2713; Fig. 1G-H, see row 1 and 3). Next, we developed a virus for which Cx35_M1_ expression is suppressed by Cre-recombinase negative (AAV-DO-Cx35_M1_-FLAG-T2A-mCherry, Fig. 1D, see also Supplemental Fig. S4 and methods). Again, we injected *Sapap3-/- ::PV-Cre* mice with CaMKII-Cx34.7_M1_ in OFC and AAV-DO-Cx35_M1_ in Str. We observed significantly lower grooming at 5 weeks, compared to mice that expressed only the fluorophores; however, no statistical differences were observed at 10 or 15 weeks (Dunn’s multiple comparisons test p=0.0228 at 5 weeks; Z=1.941, adjusted p=0.2610 at 10 weeks; Z=0.3811, adjusted p=>0.9999 at 15 weeks all for AAV-DO-Cx35_M1_ in Str; Supplemental Fig. S2B). Similarly, this LinCx targeting approach had no impact on the size of their facial lesions, compared to control mice that expressed both fluorophores (Dunn’s multiple comparisons test Z=1.429, p=0.7646; Fig. 1G-H, see row 1 and 6). Thus, our results indicated that LinCx expression across the OFC⟶Str synapses had to be targeted to PV+ interneurons to prevent pathological grooming behavior.

To test whether other Str neurons could also prevent this pathological behavior, we examined the impact of LinCx expression across OFC-striatal SPN synapses. We generated two mouse lines, enabling us to independently target D1- or D2-expressing SPNs in the knockouts: *Sapap3-/- ::D1-Cre* and *Sapap3-/- ::A2A-Cre* mice. We then injected CaMKII-Cx34.7_M1_ into OFC and DIO-Cx35_M1_ into Str of each line (Fig. 1I and 1M). Control mice were injected with CaMKII-EGFP into OFC and DIO-mCherry into Str. No significant difference in grooming time was found between fluorophore-only and LinCx-edited mice for either line (F_2,35_ = 0.09477; p=0.9098 for virus group × time effect; F_1,19_=0.03, p=0.8646 for *Sapap3-/- ::D1-Cre* mice using mixed-effects model ANOVA. F_2,36_ = 0.1323; p=0.8765 for virus group × time effect; F_1,18_=0.0267, p=0.8731 for *Sapap3-/- ::A2A-Cre* mice using mixed-effects model ANOVA, see Fig. 1J and 1N). Similarly, we observed no difference in facial lesion size 15 weeks after viral injection (U=54, p=0.95 for *Sapap3-/- ::D1-Cre* mice; U=48, p=0.95 for *Sapap3-/- ::A2A-Cre* mice using two-tailed Mann-Whitney U test; Fig. 1K-L and Fig 1O-P). Notably, independent of LinCx expression, we observed a strain effect on *Sapap3* -/- lesion size. Lesion size for the *Sapap3-/- ::D1-Cre* and *Sapap3-/- ::A2A-Cre* lines were smaller than observed in *Sapap3-/- ::PV-Cre* mice (post-hoc Kruskal-Wallis test H=9.802, p=0.0074 for comparison of lesion size across the *Sapap3*-/- fluorophore control groups for the 3 Cre lines), potentially due to a higher degree of backcrossing in the *Sapap3-/- ::PV-Cre* line. Nevertheless, together, these results showed that only selective expression of LinCx across the OFC⟶Str PV+ synapses was sufficient to prevent pathological grooming behavior.

We next tested whether LinCx editing of the OFC⟶Str PV+ circuit impacted neuronal activity in Str (Fig. 2A-B). We first characterized the impact of *Sapap3* deletion using *Sapap3*+/+ and *Sapap3*-/- mice. Mice were chronically implanted with microwires targeting OFC and a 1024-channel silicon probe targeting Str (Fig. 2B). After 2 weeks of recovery, OFC local field potential (LFP) activity and Str single-unit activity were obtained concurrently in awake freely behaving mice (Fig. 2C). We classified Str single unit activity based on their waveforms. Single units with a spike half-width between 130 and 250µs and mean firing rates less than 5Hz were classified as SPNs (Fig. 2D, left). Single units with a spike half-width between 70 and 120 were classified as narrow spiking units (NSUs, Fig. 2D, right). When we compared *Sapap3*+/+ and *Sapap3*-/- mice, we found that the *Sapap3*-/- mice exhibited a lower proportion of NSUs (χ2 = 20.64, df=2, p<0.01 using Chi-square test, Fig. 2E). In contrast to prior work^17^, we also found that the firing rate of SPNs was lower in the *Sapap3*-/- mice compared to controls (U=16482, approx. p=0.0057 using two-tailed Mann-Whitney U test, Fig. 2G, left). We observed no differences in the firing rate of the NSUs between the groups (U=2886, approx. p=0.2598 using two-tailed Mann-Whitney U test, Fig. 2G, right). Since PV+ interneurons exhibit spike waveforms consistent with NSUs^17^, we also tested whether the lower proportion of NSUs observed in *Sapap3*-/- mice was due to a loss of PV+ interneurons. When we quantified PV-expressing cell density by immunofluorescence (Fig. 2I), we found no significant difference in PV-expressing cell density between *Sapap3*+/+ and *Sapap3*-/- mice (Unpaired t-test, t=0.4289, df=9, p=0.68, n=5/6, Fig. 2J, left).

**Figure 2:**
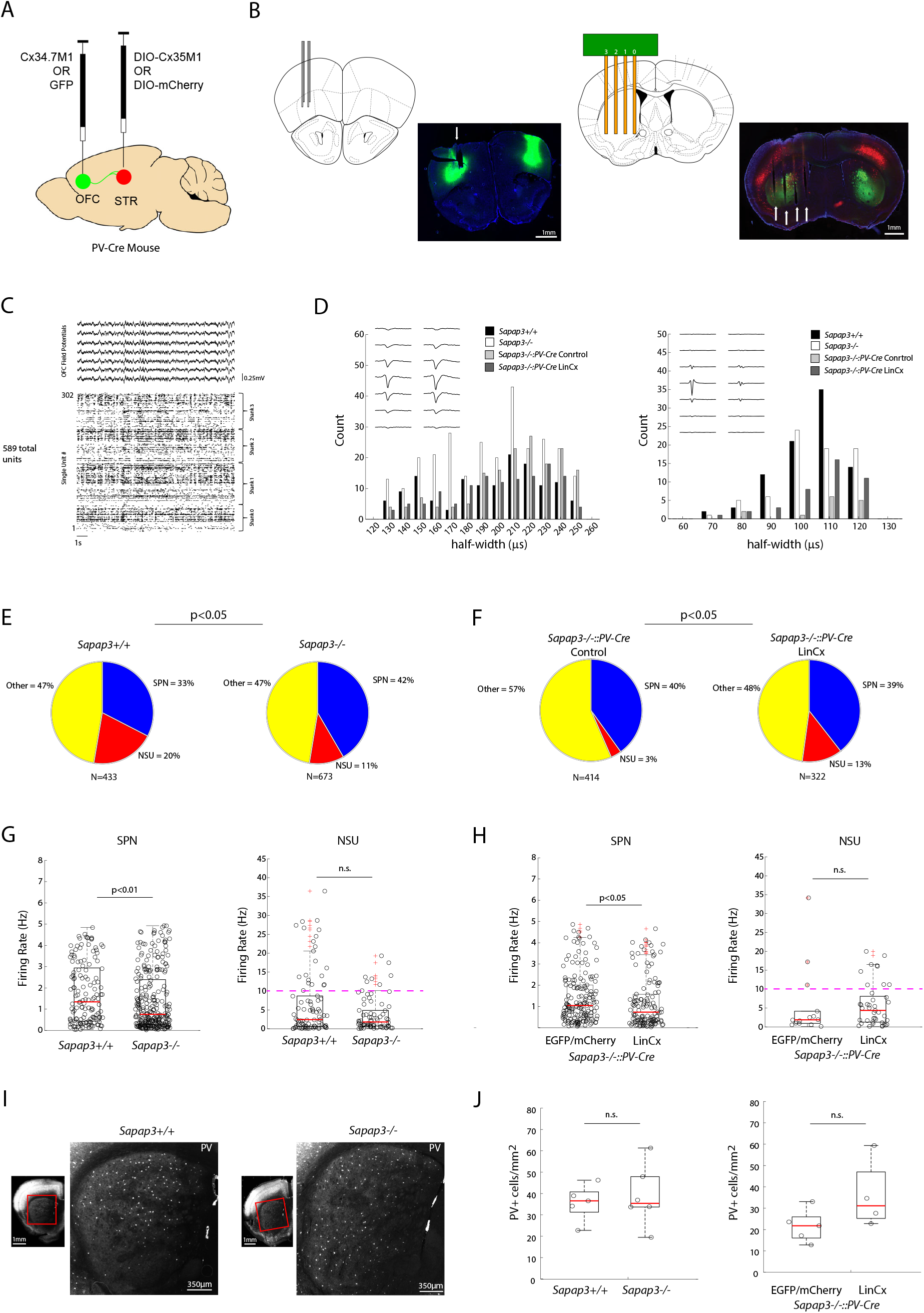
OFC⟶Str-PV circuit editing recovers striatal narrow spiking unit activity in *Sapap3*-/- mice. **A)** Viral target strategy for OFC⟶Str PV+ circuit. **B)** Schematic and representative image of OFC electrode targeting (left). Green fluorescence indicates cells expressing Cx34.7_M1_-T2A-mEmerald. Schematic and representative image of silicon probe targeting Str (right). Green fluorescence indicates terminals from OFC expressing Cx34.7_M1_-T2A-mEmerald. Red fluorescence indicates PV+ interneuron soma that expresses AAV-hSyn-DIO-Cx35_M1_-T2A-mCherry. **C)** Representative OFC LFPs and raster plot depicting activity of 302 single units recorded concurrently from central Str. **D)** Waveforms distribution for putative striatal projection neurons (SPNs, spike half-with between 130 and 250µs, and mean firing rate < 5Hz, left) and narrow spiking units (NSU, spike half-with between 70 and 120µs, right). **E)** Portion of cells characterized as SPNs or NSUs in *Sapap3*-/- vs. *Sapap3*+/+ mice (p<0.05 using Chi-square test). **F)** Portion of cells characterized as SPNs or NSUs in LinCx edited *Sapap3-/- ::PV-Cre* mice vs. non-edited EGFP/mCherry fluorophore-only controls (p<0.05 using Chi-square test). G-H) Mean firing rates of SPNs and NSUs in **G)** *Sapap3*-/- vs. *Sapap3*+/+ mice, and **H)** LinCx edited *Sapap3-/- ::PV-Cre* mice vs. non-edited controls (*p<0.05 using two wailed Mann-Whitney U test). The purple dashed line indicates the firing threshold classically utilized to characterize putative PV+ interneurons. **I)** Representative images of immunohistochemistry against PV in *Sapap3*+/+ and *Sapap3*-/-. Red boxes indicate region-enlarged insets to the right. **J)** Quantification of PV cell density in Str based on immunohistochemistry (left: t=0.4289, df=9, p=0.68 using a two-tailed unpaired t-test; right: t=1.1778, df=7, p=0.12 using a two-tailed unpaired t-test). For box and whisker plots, the central mark is the median, the edges of the box are the 25th and 75th percentiles, the whiskers extend to the most extreme datapoints the algorithm considers to be not outliers, and the outliers are plotted individually as a “+”.

After characterizing the impact of *Sapap3* deletion on Str single-unit firing rates, we tested how LinCx editing of the OFC⟶Str PV+ circuit impacted cellular firing. *Sapap3-/- ::PV-Cre* were virally infected with Cx34.7_M1_ and Cx35_M1_ at 7-8 weeks. Control mice were infected with fluorophores. Mice were then implanted with microwires targeting OFC and a silicon probe targeting Str 15-20 weeks later (Fig. 2A-B). Again, we recorded neural activity following surgical recovery, and Str single-unit activity was characterized based on the spike waveforms (Fig. 2C-D). Strikingly, LinCx OFC⟶Str PV+ circuit expression increased the proportion of NSUs observed in the *Sapap3-/- ::PV-Cre* mice relative to controls (χ2 = 24.71, df=2, p<0.01 using Chi-square test, Fig. 2F). LinCx OFC⟶Str PV+ circuit expression also decreased the firing rate of SPNs but had no impact on the firing rate of NSU between the groups (SPN: U=8925, approx. p=0.0142; NSU: U=233.5, p=0.3068, using two-tailed Mann-Whitney U test; Fig. 2H). No group differences were observed in the Str PV+ cell density (Unpaired t-test, t=1.778, df=7, p=0.12, n=5/4, Fig. 2J, right). Thus, *Sapap3* deletion suppressed Str SPN activity and LinCx OFC⟶Str PV+ circuit expression suppressed it further. On the other hand, LinCx expression in the OFC⟶Str PV+ circuit restored the activity of NSUs that were otherwise silenced by *Sapap3* deletion.

As LinCx functions by increasing the electrical coupling of neurons, we next wanted to investigate how LinCx editing of the OFC⟶Str PV+ circuit impacted the entrainment of Str neurons to OFC oscillations. We observed a peak in OFC LFP oscillatory power between 1-5Hz (Fig. 3A, top); thus, we filtered the LFPs to isolate activity in this frequency range (Fig. 3A, bottom). We then extracted the phase of the filtered OFC LFPs and quantified the relationship between single unit activity at this phasic activity (Fig. 3B-C). Finally, we introduced temporal offsets ranging between -200ms and 200ms between the single-unit activity and the LFP phase, enabling us to also quantify the relationship between Str single unit activity and OFC activity in the immediate past and immediate future^23,24^. Indeed, we found that when we pooled SPN and NSU single units in *Sapap3*+/+ mice, Str neurons optimally coupled to OFC activity in the immediate past or immediate future (Fig. 3D). In contrast, *Sapap3* deletion increased coupling to OFC 1-5Hz activity across all temporal offsets (Fig. 3E). Interestingly, when we limited our analysis solely to SPNs, we still observed increased coupling to OFC oscillations ranging from - 200ms in the past to 40ms in the future in the *Sapap3*-/- mice (Supplemental Fig. S5A). Thus, the increased coupling observed in the *Sapap3*-/- mice did not simply reflect the decreased portion of NSUs represented by their population activity. Finally, we repeated this analysis in the *Sapap3-/- ::PV-Cre* that had been treated with LinCx editing of the OFC⟶Str PV+ circuit, as well as control mice which solely expressed fluorophores. Strikingly, LinCx editing decreased the coupling of Str neuronal activity to OFC activity in the immediate past and immediate future (Fig. 3F). Similar results were observed when we limited our analysis to SPNs (Supplemental Fig. S5B). Thus, *Sapap3* deletion increased the coupling of SPNs to OFC activity, while LinCx editing of the OFC⟶Str PV+ circuit decreased this coupling in a temporally selective manner. Together, these findings established the OFC⟶Str PV+ circuit in mediating pathological grooming in the *Sapap3*-/- mice.

**Figure 3:**
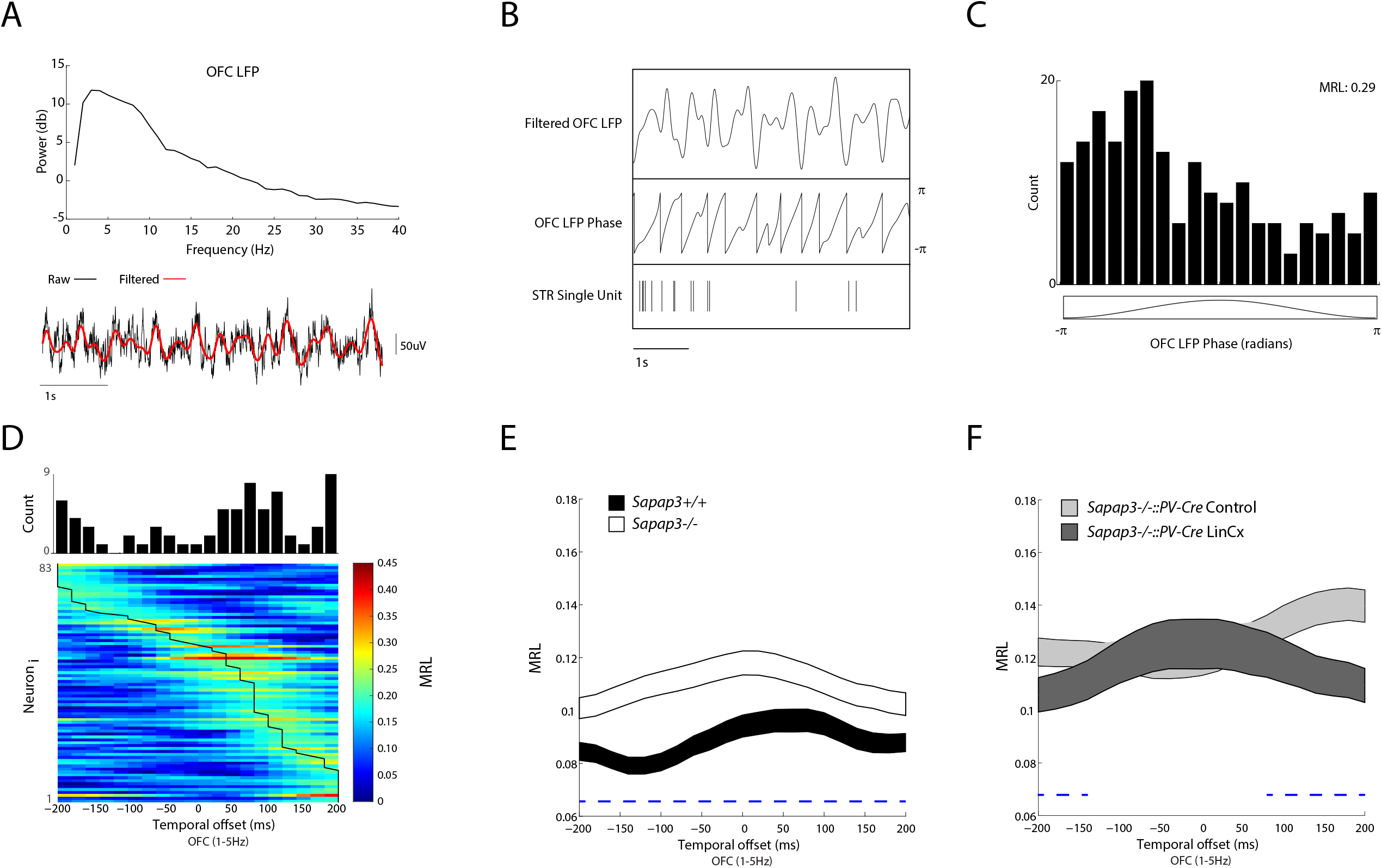
LinCx circuit editing moderates OFC⟶Str spike timing deficits in *Sapap3*-/- mice. **A)** Representative power spectral density for OFC LFPs recorded from a *Sapap3*+/+ mouse. Note the peak in activity between 1 and 5Hz (top). Representative OFC LFP with the 1-5Hz filtered LFP signal shown in red (bottom). **B)** Plot showing a 1-5Hz filtered OFC LFP (top), the extracted oscillatory phase (middle), and a concurrently recorded Str single unit (bottom). **C)** The firing of the same Str unit relative to the phase of the OFC 1-5Hz oscillation. **D)** Phase locking of Str units recorded in *Sapap3*+/+ mice. Heatmap depicts the mean resultant length (MRL) of the 83/433 units that phase-locked to OFC oscillations with temporal shifts ranging from -200 to 200ms. The black line highlights the optimal temporal offset for each phase-locked cell. The histogram above depicts the distribution of peak temporal offsets for the 83 neurons. Significance was determined using the Rayleigh test at α=0.05/21 temporal shifts. **E-F)** Coupling of pooled Str SPN and NSU units to OFC 1-5Hz oscillations at temporal offset ranging from -200 to 200ms. E) Str phase-coupling in *Sapap3*-/- vs. *Sapap3*+/+ mice and F) LinCx-edited *Sapap3-/- ::PV-Cre* mice vs. non-edited controls. The dashed blue line highlights offsets at which significant differences were observed between groups using a Mann-Whitney U test with a false discovery rate correction for comparisons across the 21 offsets. Data shown as mean ± s.e.m.

## Discussion

Here we set out to test whether circuit-selective assembly of an electrical synapse could be utilized to edit a genetically mediated chemical synaptic disruption and restore normal behavior. Using the *Sapap3*-/- mutant mouse model, we found that LinCx editing of the OFC⟶Str PV+ circuit corrected spike timing between OFC and Str, and it ameliorated the mutant animals’ pathological repetitive behavior. Such a behavioral outcome was observed when the striatal portion of LinCx synapse was restricted to PV+ interneurons, highlighting the specificity of the editing approach (Fig. 4). Previous in vivo work has implicated hyperactivity of striatal SPNs in mediating the repetitive grooming displayed by *Sapap3*-/- mice^17^. On the contrary, here we observed a decreased SPN firing in *Sapap3*-/- mice compared to controls. Our findings were consistent with the reduced field excitatory postsynaptic potentials described in *Sapap3*-/- mice^7^. Prior in vivo work also implicated hypoactivity of striatal PV+ interneuron activity^17^. Though we observed a lower portion of NSUs in *Sapap3*-/- mice using our high-density striatal recordings, *Sapap3*-/- mice did not exhibit a lower NSU firing rate. Interestingly, LinCx editing further lowered the firing rate of striatal SPNs and had no impact on the firing rate of striatal NSUs. Rather, LinCx editing increased the proportion of NSUs we observed in the *Sapap3-/- ::PV-Cre* mice. Since we observed a similar density of PV+ interneurons compared to controls in our histological analysis, our findings suggest that an overall physiological silencing of Str NSUs may be central to the repetitive grooming phenotype that results from *Sapap3* deletion. Striatal SPNs also showed increased coupling to LFP activity in OFC in the *Sapap3*-/- mice. Such coupling likely reflects a loss of feedforward inhibition in the OFC⟶Str circuit due to the silencing of NSUs within central Str. LinCx editing of the OFC⟶Str PV+ circuit both ameliorated the pathological grooming behavior and reduced the increased coupling in the *Sapap3* mutant mice. These findings raise the possibility that altered SPN spike timing across the OFC⟶Str circuit, rather than simply changes in Str SPN firing rates alone may serve as a core pathological mechanism underlying the repetitive behavioral phenotype in *Sapap3*-/- mice. OCD impacts 1-3% of the global population^25^. Yet, standard treatments including cognitive behavioral therapy and SSRI-based pharmacotherapy are only effective in a subset of individuals with OCD^26-28^. Furthermore, these pharmacotherapeutics require chronic administration, with symptoms recurring in the setting of medication non-compliance. Highlighting the predictive validity of the *Sapap3*-/- mouse model for studying OCD, chronic SSRI-based treatment also ameliorates this model’s repetitive behavior^7^. Interestingly, anxiety is highly co-morbid with OCD, and it has been reported that *Sapap3*-/- mice have exhibited anxiety-like behavior^7^. Further supporting the predictive validity of the *Sapap3*-/- model, their anxiety state is also reduced by chronic SSRI treatment^7^. While we also confirmed an anxiety-like phenotype in *Sapap3*-/- mice (Supplemental Fig. S6), we found that editing of OFC⟶Str PV+ circuitry had no impact on this behavioral phenotype. Thus, though LinCx precision editing of the OFC⟶Str PV+ circuit yielded amelioration of repetitive behavior, it had no impact on another major psychiatric phenotype of the model. Overall, this finding suggests that the therapeutic impact of chronic SSRI-based treatment on anxiety in OCD may be mediated at other distinct circuits in the brain, either via its direct effects on transporters or via secondary effects whereby released serotonin may interact directly with DNA in a manner that subsequently changes gene expression^29^. Future studies in which LinCx is used to edit other chemical synapses in the *Sapap3*-/- brain may help to clarify whether other circuits can be targeted to potentially reduce or prevent the development of anxiety.

**Figure 4:**
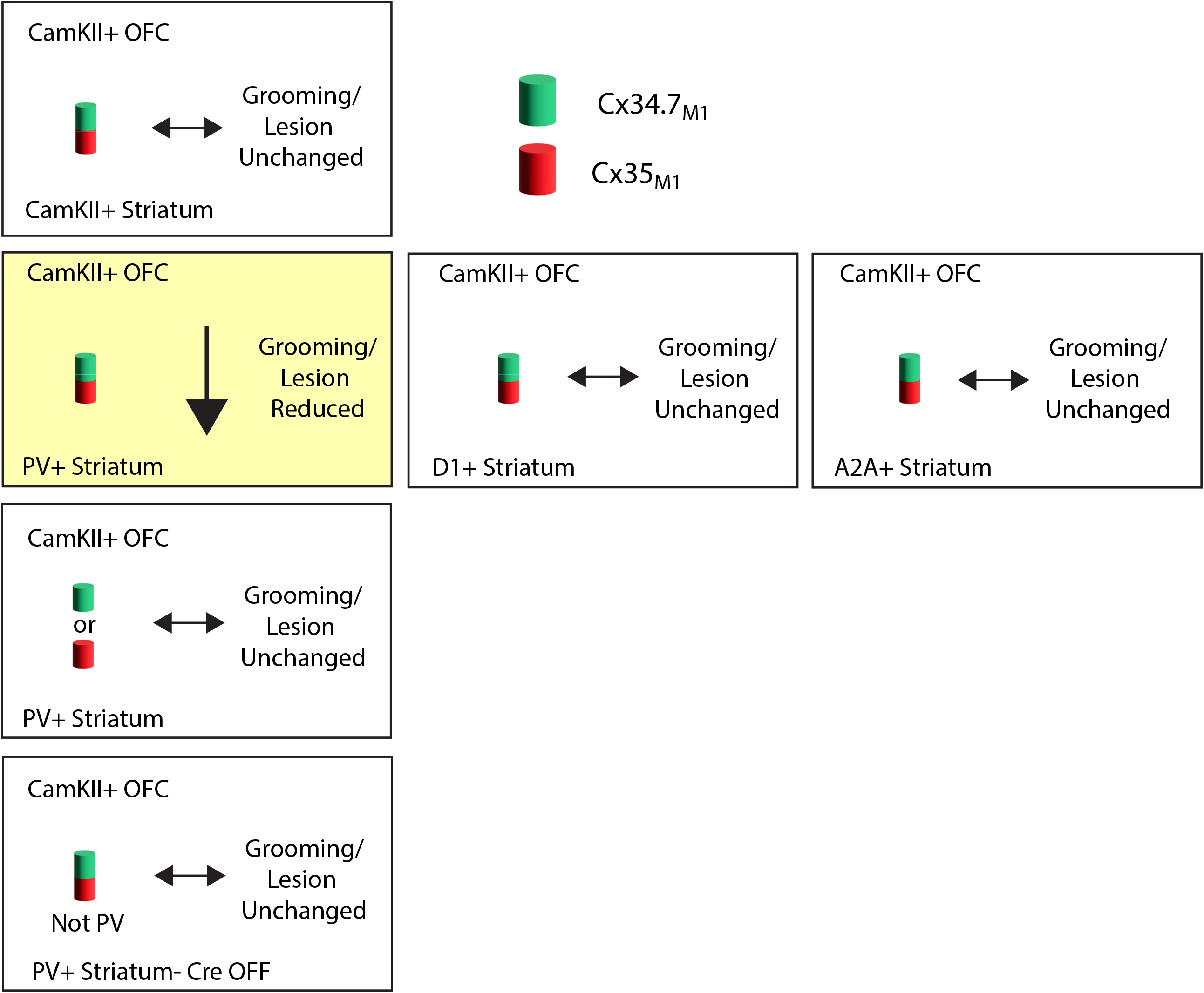
Summary of LinCx precision circuit editing manipulations and corresponding effects on repetitive behavior in *Sapap3* deficient mice. Expression of Cx34.7_M1_ (green cylinder) and Cx35_M1_ (red cylinder) in cell type-specific OFC⟶Str circuits, with arrows indicating alterations (if any) to grooming and lesion formation.

Notably, virally mediated expression of the *Sapap3* protein in juvenile *Sapap3*-/- mice prevents the emergence of repetitive grooming and anxiety-like behavior in adulthood^7^. This directly argues that deficient striatal *Sapap3* mediates the behavioral dysfunction displayed by *Sapap3*-/- mice. Moreover, this prior work broadly implicates striatal-dependent circuits in mediating repetitive behavior and anxiety-like phenotypes in OCD. Yet this approach of expressing *Sapap3* in human Str during early development is highly unlikely to translate to a clinically viable therapeutic, since most individuals with OCD do not have a genetically mediated disruption of *Sapap3*. Rather, an approach that targets a convergent circuit-based phenotype shared across the many different genetic mediators of OCD bears greater potential for preventative therapy. Here, we demonstrate that precision circuit editing using an electrical synapse can override a genetically mediated synaptic deficit and prevent the emergence of behavioral disruption relevant for OCD and other neuropsychiatric disorders in adulthood. Future work will be necessary to test whether such a circuit-editing framework extends to models of other genetic lesions linked to OCD, and to models of other psychiatric disorders.

## Acknowledgments

We would like to thank Dr. Elizabeth Ransey for co-designing and cloning 2 viral constructs utilized for the first time in this study, Nicholas B Clark for assistance with histology, animal husbandry, and for cloning a viral construct, Theodore J. Moser for assistance with animal husbandry, Adam Weissman for assistance with analyzing behavioral data, Erica Robinson for performing preliminary experiments, Dr. Jean Mary Zarate for revising the manuscript, and Dr. Stephen D. Mague and Arielle Ramey for administrative support for the scientific program. We would also like to thank the Duke Viral Vector Core for producing LinCx viruses used in this study, the Duke Light Microscopy Core for confocal microscopy support, and the UCI Center for Neural Circuit Mapping Viral Core for pseudorabies and the associated helper virus. This work was supported by the Hartwell Postdoctoral Fellowship to K.W.C.; NIH Grants R01NS110059 and R01NS133430 to N.C.; and Duke University School of Medicine MedX Grant, Duke University Chancellor’s Discovery award, Duke University School of Medicine Kahn Neurotechnology Grant, Hope for Depression Research Grant, and NIH Grants R21EY029451, R01MH120158, R01MH125430 and 1DP1MH132709 to KD. A special thanks to Freeman A. Hrabowski, Robert and Jane Meyerhoff, and the Meyerhoff Scholarship Program from KD.

## Author contributions

Conceptualization and Methodology – K.W.C., N.C., and K.D.; Formal Analysis – K.W.C., H.A.S., and K.D.; Investigation – K.W.C., and H.A.S., K.D.; Resources – N.C., and K.D.; Writing – Original Draft – K.W.C., and K.D.; Writing – Review & Editing – K.W.C, N.C., and K.D.; Visualization – K.W.C, and K.D.; Supervision – K.W.C, and K.D.; Project Administration and Funding Acquisition – K.W.C, N.C., and K.D.;

See Supplemental materials for detailed author contributions.

## Methods

### Animal Care and Use

All experimental procedures were performed in compliance with approved protocols from the Duke University Institution of Animal Care and Use Committee at Duke University. *Sapap3*-/- mice and littermate controls were bred from heterozygous breeding pairs. *Sapap3-/- ::Cre* recombinase mice were generated by crossing *Sapap3*-/- and +/-mice to respective Cre+ mice: PV+ interneurons targeted using B6 Pvalb-IRES-Cre (Jackson Laboratory #017320); D1+ striatal projection neurons targeted using D1-Cre (Jackson Laboratory #037156); and D2+ striatal projection neurons targeted using Adora2a-Cre (gift from Dr. Henry Yin; MMRC 34744). *Sapap3*-/- ::Cre+ mice were crossed together to generate *Sapap3*-/- ::Cre+ offspring used for experiments. Mice were genotyped prior to weaning using standard protocols for Cre and *Sapap3*-(Transnetyx). 2-5 mice were housed together by sex and age. Mice were given unrestricted access to food and water and were maintained on a 12-hour light cycle. All experiments were conducted during the light-on phase of their light cycle. Humane endpoints were defined as follows: animals that exhibited a 20% weight loss over a week, active bleeding for more than 5 minutes, or bilateral lesions over 5mm in size. 3 mice from the *Sapap3-/- ::PV-Cre* mice virally transduced with Cx34.7_M1_-only group met criteria for euthanasia prior to the completion of the 15-week behavioral experimental protocol.

### Molecular Cloning and Viral Production

pAAV-CaMKIIa-Cx34.7_M1_-2xHA-T2A-mEmerald was generated by producing a gBlock with Cx34.7_M1_-2xHA-T2A-mEmerald (Integrated DNA Technologies) and ligating together with BamHI-HF and EcoRI-HF digested pAAV-CaMKIIa-EGFP (Addgene # 50469, a gift from Bryan Roth). Similarly, pAAV-CaMKIIa-Cx35_M1_-FLAG-T2A-mCherry was generated likewise but using a gblock containing Cx35_M1_-FLAG-T2A-mCherry. pAAV-hSyn-DO-Cx35_M1_-FLAG-T2A-mCherry(off)-iRFP670(on)-WPRE was generated by amplifying Cx35_M1_-FLAG-T2A-mCherry using AscI and NheI and iRFP670 with NheI and ligating step-wise into pAAV-hSyn-DIO-mCAR(off)-CheTA(on)-WPRE (Addgene # 11387, a gift from Adam Kepecs^30^).

Phusion using GC Phusion buffer under standard conditions was used to amplify DNA fragments. All restriction enzymes were from NEB. Ligations were completed with standard In-Fusion protocol (Takara Bio). Plasmids were transformed into Stbl3 competent E. coli (ThermoFisher). Final vectors were full length sequenced prior to viral production (Eton Bio).

All Cx viruses were AAV2/9 serotype and produced at the Duke Viral Vector Core (graded for in vivo use).

### Surgery

#### Viral Manipulation Surgery

Seven-week-old mice were anesthetized with isoflurane (1%) and then secured in a stereotaxic device. A blunt end 33-gauge Hamilton syringe was used to deliver 100nL of AAV9-CaMKII-Cx34.7_M1_ -T2A-mEmerald (5 × 10^13^ vg/ml) or AAV9-CaMKII-EGFP (Addgene viral prep # 50469-AAV9, titer = 2.8 × 10^13^ vg/ml, a gift from Bryan Roth) bilaterally, into orbital frontal cortex (OFC; +2.6mm AP and ±2.0mm ML from bregma, and -1.49DV from brain surface at a 10° angle) and 250nL AAV9-hSyn-DIO-Cx35_M1_-T2A-mCherry (1.26 × 10^13^ vg/ml)^22^, AAV9-hSyn-DIO-mCherry (Addgene viral prep # 50459-AAV9, titer = 2.3 × 10^13^ vg/ml, a gift from Bryan Roth), AAV9-CaMKII-Cx35_M1_ -T2A-mCherry (6.24 × 10^13^ vg/ml), or AAV9-hSyn-DO-Cx_M1_ 35 -T2A-mCherry (3.5 × 10^12^ vg/ml) into striatum (Str; +0.3mm AP and ±2.35mm ML from bregma, and -2.35mm DV from brain surface at a 10° angle). Virus was delivered at a rate of 100nL/min using a microinjection syringe pump (WPI Micro2T SMARTouch with UMP3 pumps). Carprofen (5mg/kg, s.c.) injections were administered once prior to surgery and again every 24 hours for three days following viral injection surgery.

#### Viral Tracing Surgery

At 22 weeks, mice were anesthetized with isoflurane (1%) and secured in a stereotaxic device. A blunt end 33-gauge Hamilton syringe was used to deliver 250nL of AAV8-DIO-hSyn-TC66T-2A-mRuby3-2A-OG (2.16 × 10^10^ vg/ml, UCI Center for Neural Circuit Mapping Viral Core) into Str (+0.3mm AP and ±2.35mm ML from bregma, and -2.35mm DV from brain surface at a 10° angle) using a microinjection syringe pump (WPI Micro2T SMARTouch with UMP3 pumps) at a rate of 100nL/min. After 21 days, mice were then injected with 250nL EnvA-SAD-B19-RVdG-mNeonGreen (5 × 10^7^ vg/mL, UCI Center for Neural Circuit Mapping Viral Core) as with the same conditions and location as the AAV8 delivery method previously reported ^31^. Carprofen (5mg/kg, s.c.) injections were administered once prior to each surgery and again every 24 hours for three days following each viral injection surgery.

#### Electrode Implantation and Opsin Viral Surgery

After 15 weeks post-viral injection, a subset of mice was anesthetized with isoflurane (1%), placed in a stereotaxic device, and metal ground screws were secured to the anterior cranium and posterior cortex. A single ground wire was connected to the front screw and to one of the posterior screws. As before, viral injections were performed using a blunt end 33-gauge Hamilton syringe to deliver 150nL of AAV1-CAG-fDIO-ChR2-mCherry (Addgene viral prep # 75470-AAV1, titer = 2.3 × 10^13^ vg/ml, a gift from Connie Cepko^32^) unilaterally into OFC (+2.6mm AP and -2.0mm ML from bregma, and -1.49mm DV from brain surface at a 10° angle) and 250nL rAAV-EF1a-FLPo (Addgene viral prep # 55637-AAVrg, titer = 1.7 × 10^13^ vg/ml, a gift from Karl Deisseroth^33^) unilaterally into Str (+0.3mm AP and -2.35mm ML from bregma, and -2.35DV from brain surface at a 10° angle). Virus was delivered at a rate of 100nL/min using a microinjection syringe pump (WPI Micro2T SMARTouch with UMP3 pumps). Following viral infusion, a custom 8-microwire bundle (built in a 0.25mm spaced square grid using 2×2 arrangement with 2 wires in each hole with 0.2mm DV stagger for the wires in the same hole) with an 100µm optic fiber in the center (MFC_100/1.5-0.22_8mm_MF2.5_FLT, Doric Lenses) connected to a 32-pin omnetics connector was inserted into OFC (+2.5mm AP, -1.5mm ML, -1.7mm DV) and secured with dental cement to the anterior screw. After drying, a 1024-channel silicon probe (NeuroNexus, SINAPS_4S_1024) was inserted into Str (+0.5mm AP, -3mm through -1mm ML based on placement of medial probe at -1.0mm ML from bregma, and -4mm DV from the brain surface). Dental cement was liberally applied to secure the probe to the skull. Dental cement and rubberized glue were used to secure the implants in place. Again, carprofen (5mg/kg, s.c.) injections were administered once prior to surgery along with 0.1ml 10% dextrose and 0.6ml saline and again every 24 hours for three days following viral injection surgery.

### Behavior

Grooming behavior was quantified in *Sapap3*-/- and control mice at age 23 weeks and anxiety behaviors were quantified at 17 weeks. Grooming behavior was quantified in virally injected mice at 5, 10, and 15 weeks post-surgery (ages 13, 18, 23 weeks). Anxiety-related behavior was assessed in the same animals 10 weeks post-surgery (∼age 17 weeks) as described below.

#### Open Field Test for Grooming

Mice were habituated to handling for 3 days prior to experimentation and placed into the experimental room at least 1 hour prior to behavioral testing. Lighting was set for 50lux inside of either a black or white open field box (45cm x 45cm x 30cm) based on coat color to provide maximal contrast for tracking purposes. Mice were placed into the box and allowed to explore freely for 5 min (baseline). Immediately after, mice were misted with a single spray of tap water (∼300µL) to induce grooming (i.e., water mist-induced grooming). This data was not utilized in this study. Videos were recorded for an additional 10 min. Experimental videos were analyzed using Ethovision XT17 with mouse behavior recognition module (Noldus). Grooming was identified by the Ethovision software with threshold settings based on >90% agreement with a blinded observer for 2 videos for each set of experimental settings (arena and detection settings). Behavioral videos that were not able to be tracked with Ethovision were manually scored by a blinded observer (n=6).

#### Bright Open Field Test

The bright open field test was conducted as before^34^. In brief, mice were habituated to the behavioral room at least 1 hour prior to behavioral testing. Lighting was set for 500lux at the center of the open field box (45cm x 45cm x 30cm). Mice were placed into the periphery of the box and allowed to freely explore for 5min. Mouse position was tracked using Ethovision XT17 (Noldus) and center time was calculated based on presence of the body center in the innermost third of the arena.

#### Light/Dark Emergence Test

The light dark emergence test was conducted in a rectangular arena (60cm x 20cm x 20cm), where a small portion (20cm x 20cm) was covered with a dark plastic lid that allowed for IR imaging and was closed by a small door covering an opening (5cm x 5cm) (Noldus). Room light was set to 500lux. Mice were placed in the room for at least 1 hour prior to testing. During testing, mice were placed into the dark side of the chamber. The door was removed and mice were allowed to freely explore for 5 min. Videos were tracked using Ethovision XT17 (Noldus) and time in light side of the chamber was measured based on center of the mouse. Videos that were not able to be tracked in Ethovision were manually scored for time in light chamber by a blinded observer (n=16).

#### Elevated Plus Maze

The elevated plus maze test was conducted as previously described^34^. In brief, we used a cross-shaped arena comprised of four arms wherein 2 opposing arms are enclosed by walls 16.5cm high (closed arm), and the other two arms are opened with a 1mm rim (open arm). All arms were 30cm long and 5cm wide, connected by a center region measuring 5cm x 5cm. The arena was elevated 91.4cm above the floor, and lighting at the ends of the open arms was set to 175lux. Mice were placed into the center of the arena with their nose in a closed arm, and they were allowed to freely explore for 5min. Video recordings were tracked on Ethovision XT17 (Noldus). Duration in the open arm was calculated based on presence of the nose in either open arm.

### Facial Lesion Scoring

Facial lesions were assessed at 24 weeks of age or earlier for mice that reached a humane endpoint (n=3). Mice were anesthetized and photographed on profile from both sides. Lesions, identified by red exposed skin, were measured and summed for each side of the face using a standard ruler and rounded to the nearest millimeter. Lesion scoring was based on the average size of lesions on the left and right side of the face (i.e., total mm of lesions/2).

### Acquisition and Analysis of Neural Activity

Mice were habituated to the recording room in a new homecage for at least 60 minutes over multiple days prior to testing. On the recording day, mice were connected to a headstage (Blackrock Microsystems, UT, USA) without anesthesia and placed in the home cage used for habituation. Mice were also connected to a mezzanine board and SiNAPS interface cable (Neuronexus, MI USA). Mice were habituated to the recording room for at least 60 minutes on the testing day. For each mouse, the last 10 minutes of neural data, which corresponded to low activity periods, were used for subsequent electrophysiological analysis. We selected these periods since our prior work demonstrated that phase coupling is diminished during novelty exposure and exploration^35,36^.

Neuronal activity was sampled from OFC at 30kHz using the Cerebus acquisition system (Blackrock Microsystems Inc., UT). Local field potentials (LFPs) were bandpass filtered at 0.5– 250Hz and stored at 1000Hz. An online noise cancellation algorithm was applied to reduce 60Hz artifact. Neural data was also sampled at 20kHz using a SmartBox Pro acquisition system (NeuroNexus, MI USA). All neurophysiological recordings were referenced to a ground wire connected to both ground screws. Finally, a TTL pulse was generated using the digital output from the Cerebus acquisition system. This signal was stored at 10Hz using the analog input on the Cerebus acquisition system and at 20Hz using analog input on the acquisition on the SmartBox Pro acquisition system.

To extract single-unit activity, neural activity was converted to Neurodata Without Borders (NWB) format, high-pass filtered at 300Hz, and automatically sorted with Kilisort4^37^. Single units were identified based on ISI violations < 0.5, presence ratio > 0.9, an amplitude cutoff < 0.1^38,39^, a peak-valley ratio < 1.1. Single striatal projection neurons (SPN) neurons were classified by mean firing rate less than 5Hz and spike-width (at half-peak amplitude) within a range of 130– 250µs^17^. Narrow spiking units (NSUs) were classified by a spike width within a range of 70– 120µs^17^. Only single units that mapped to the central striatum targeted channels were used for analysis.

We solely utilized neuronal activity that occurred within the last 10 minutes of the recording for analysis. This approach controlled for differences in recording length across animals. LFP activity was averaged across all the OFC microwires and then filtered using a 3^rd^-order Butterworth bandpass filter to isolate 1-5Hz oscillations. The instantaneous phase of the filtered-averaged LFP was then determined using the Hilbert transform, and phase locking was calculated using the MATLAB (MathWorks) circular statistics tool box (i.e., the ‘circ_r’ function for the mean resultant length and the ‘circ_rtest’ function for the Rayleigh test of uniformity analyses). We also calculated phase locking after introducing temporal shits ranging from -100ms to 20ms between the OFC oscillation and the striatal spikes. Notably, only the last 200 spiking events were used to determine the phase locking of each neuron. Neurons that fired less than 200 times within the 10-minute recording experiment were excluded from further analysis. This approach ensured that any putative difference in firing rates would not skew our quantification and analysis of phase locking. For these analyses, the TTL signal was used to synchronize the Cerebus and SmartBox Pro acquisition systems.

Mice were also subjected to optogenetic stimulation of OFC neurons that project to Str following the initial neural recordings. These data were not utilized as part of this study.

### Histology

At the end of experiments, mice were fully anesthetized using 150mg/kg ketamine, 12mg/kg xylazine, and 3mg/kg acepromazine in saline and then perfused transcardially with PBS followed by 4% paraformaldehyde (PFA, EM Sciences) in 1xPBS. Brains were postfixed overnight at room temperature in 4% PFA in PBS and then transferred to 1x PBS + 0.01% Sodium azide and stored at 4°C until sectioning. Brains were sectioned coronally using a vibratome (Leica) at 40um thickness and mounted onto charged glass slides with Mowiol mounting media [24% (w/v) glycerol, 9.6% (w/v) Mowiol 4-88, 0.1M Tris-HCl pH 8.5] with 1µg/ml Hoechet 33342 and coverslipped. Images were obtained on a slide scanner microscope (Olympus VS200) using 4x or 10x objective. Confocal images were obtained on a confocal microscope (Zeiss 780 inverted) using 63x objective (accessed through Duke LMCF). Confocal z-stack images were rendered using Imaris 9.3.1.

#### HImmunohistochemistry

After tissue was sectioned, slices were placed 1xPBS + 0.01% sodium azide (PBS+azide). All tissue incubation steps were conducted at room temperature with rotational agitation unless otherwise noted. Tissue was briefly permeabilized by placing in 0.5% TritonX-100 in PBS+azide for 15 minutes. Tissue was then washed in PBS+azide and placed into blocking buffer (5% goat serum in PBS+azide+0.1%Tween-20) for three hours. Tissue was then placed into the diluted primary antibody [1:1000 Rabbit α-PV (Abcam ab11427)] in blocking buffer overnight at 4°C. The following day, tissue was washed in PBS+azide and incubated in secondary antibody [1:1000 Goat α-Rabbit IgG-AlexaFluor Plus 647 (ThermoScientific A32733TR and 1µg/ml Hoechet 33342 or 1:1000 Goat α-Rabbit IgG-AlexaFluor 405 (ThermoScientific A-31556)] diluted with blocking buffer for 2 hours. Following secondary incubation, tissue was washed with PBS+azide and mounted onto charged glass slides with Mowiol mounting media and coverslipped.

Micrograph images were obtained on a slide scanner microscope (Olympus VS200) using 4x or 10x objective. Confocal images were obtained on a confocal microscope (Zeiss 780 inverted) using 63x objective (accessed through Duke LMCF). Confocal z-stack images were rendered using Imaris 9.3.1.

For viral tracing studies, mice were perfused nine days after the injection of pseudorabies, and brain tissue was collected as described above. Brains were then sliced, mounted, and imaged on the slide scanning microscope using a 4x objective.

### Statistics

All Statistical tests were conducted using Graph Pad Prism (v10.1.1) or Matlab (R2021b). For unpaired data between two groups with non-Gaussian distributions, a two-tailed Mann-Whitney rank sum test was used. For paired data with non-Gaussian distributions and more than two groups, a mixed effects model (REML) was fit, and posthoc comparisons with Dunn’s multiple comparison tests were conducted accounting for multiple hypothesis testing using Dunnett’s method. For unpaired data with non-Gaussian distributions among more than two groups, a Kruskal-Wallis test was performed with Dunn’s multiple comparison tests as post-hoc tests with Dunnett’s method for multiple hypothesis testing correction. Relative proportions with small sample size (facial lesions) were tested with Fisher’s exact test. Relative proportions with large sample size (single units) were analyzed via Chi-square tests. Normally distributed, unpaired data between two groups with similar variance was analyzed using an unpaired, two-tailed t-test. Normally distributed, unpaired data between two groups with unequal variance was analyzed using an unpaired, two-tailed t-test with Welch’s correction. Normally distributed, unpaired data between more than two groups with unequal variance was analyzed using a Welch’s ANOVA test with post-hoc Dunnett’s T3 multiple comparisons test and p-value thresholds were adjusted using a Bonferroni correction for multiple hypotheses testing.

