## Supplemental Materials for "Synaptic editing of frontostriatal circuitry prevents excessive grooming in SAPAP3-deficient mice"

**Supplemental Table S1, Detailed Author Contributions**

| | |
|---|---|
| Kathryn Katsue Walder-Christensen | Conceived all aspects of the study and experimental plan with K.D.; co-designed all Connexin-T2A viral constructs, generated all mouse cross strains, and validated all viral constructs; conducted all viral surgeries, performed all behavioral testing and lesion scoring, performed behavioral analyses with H.A.S.; processed all tissue for histological confirmation of viral expression with H.A.S. and N.C., processed all tissue for histological confirmation of all electrophysiological targeting, and performed all immunohistochemical and tracing experiments and analysis; constructed all OFC targeting electrodes, performed electrophysiological implant surgeries, and collected electrophysiological data with K.D.; assisted with statistical testing of physiology data to K.D.; wrote and revised the original draft with K.D.; prepared figures with K.D.; oversaw technical support; and acquired funding to support the study. |
| Hannah A Soliman | Performed histological processing of brain tissue in conjunction with K.W.C.; Organized, maintained, and analyzed behavioral data in conjunction with K.W.C.; acquired lesion images in conjunction with K.W.C. |
| Nicole Calakos | Jointly conceived of the LinCx editing study (behavioral experiments) with K.W.C. and K.D.; jointly revised the first manuscript draft with K.W.C. and K.D.; provided Sapap3-/- mice to establish the colony utilized in this study and acquired funding to support the study with K.D. |
| Kafui Dzirasa | Jointly conceived the LinCx editing study (behavioral) with K.W.C and N.C.; Conceived the electrophysiological studies with K.W.C.; Performed electrophysiological studies with K.W.C.; analyzed and interpreted electrophysiological data; wrote the first draft of the manuscript with K.W.C.; revised the first draft of the manuscript with K.W.C., and N.C.; prepared and revised figures with K.W.C, supervised all aspects of the study; and acquired funding to support the study with K.W.C. and N.C. |
| **Attribution Process | Each team member outlined their individual contributions across a standard set of domains (conceptualization and methodology, formal analysis, investigation, resources, writing -original draft, writing -review & editing, visualization, Supervision, and Project Administration and Funding Acquisition), and subsequently had the opportunity to edit a summarized attribution description to their satisfaction. Contribution summaries were then shared across all team members. Each team member had the opportunity to raise concerns with regards to any other team member's outlined contributions, and issues that remained unaddressed after additional revisions were subjected to a mediation process led by the lead principal investigator. The assigned authorships and these detailed author contribution descriptions reflect the outcome of this process. |

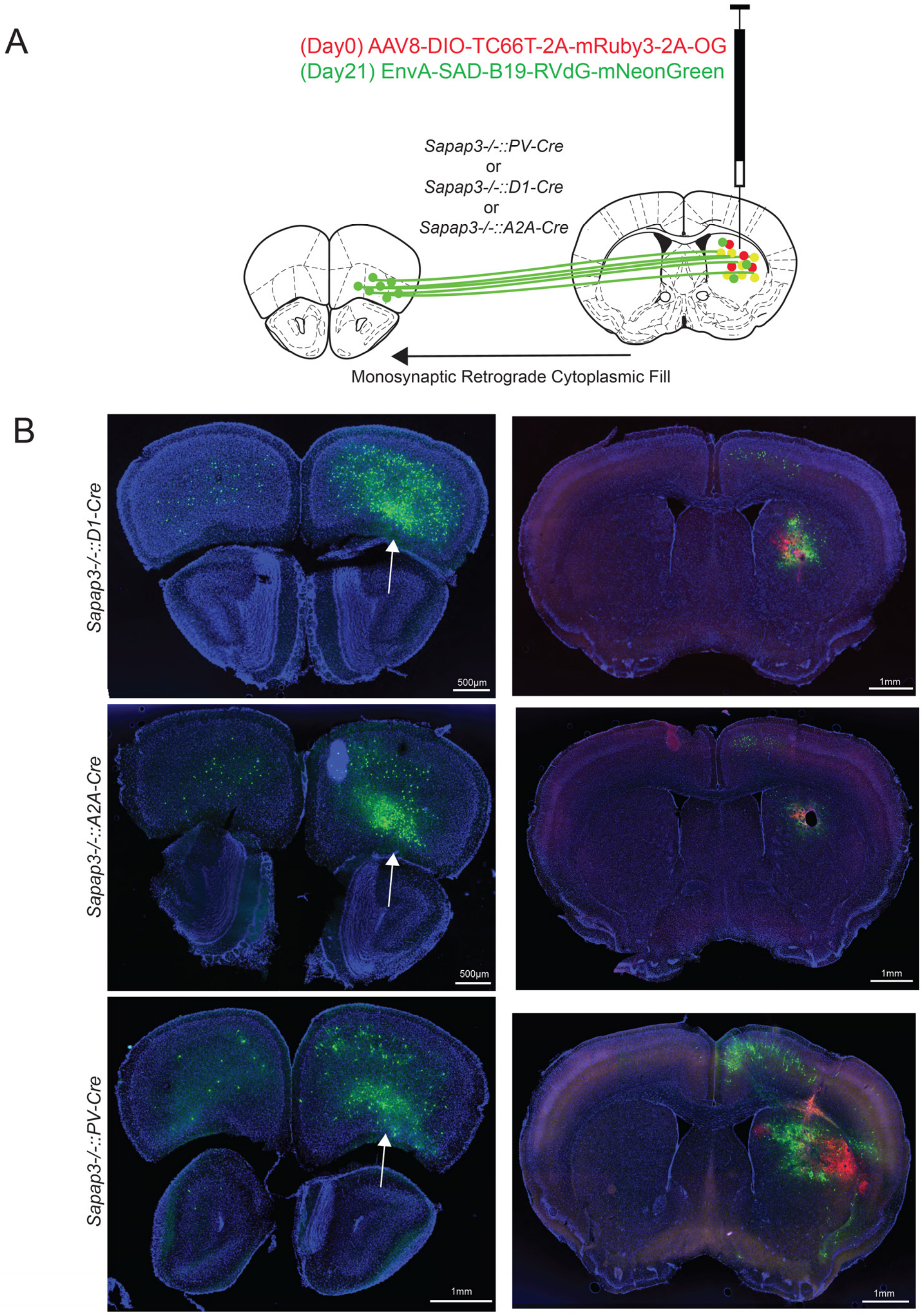


**Supplemental Figure S1: Cre-dependent retrograde labeling of OFC from D1+, A2A+, and PV+ striatal neurons in *Sapap3-/-* mice. A)** Scheme of retroviral labeling strategy for identification of innervating neurons of 3 cell populations in Str. **B)** Representative images of OFC (left) and Str (right) for *Sapap3-/-::D1-Cre, A2A-Cre, or PV-Cre* mice. Red viral expression from AAV8-DIO-TC66T-2A-mRuby3-2A-OG identifies neurons which express Cre-recombinase in Str. Green viral expression from EnvA-SAD_B19-RVdG-mNeonGreen identifies neurons that synapse onto red neurons in Str. White arrows indicate green somatic labeling in OFC.

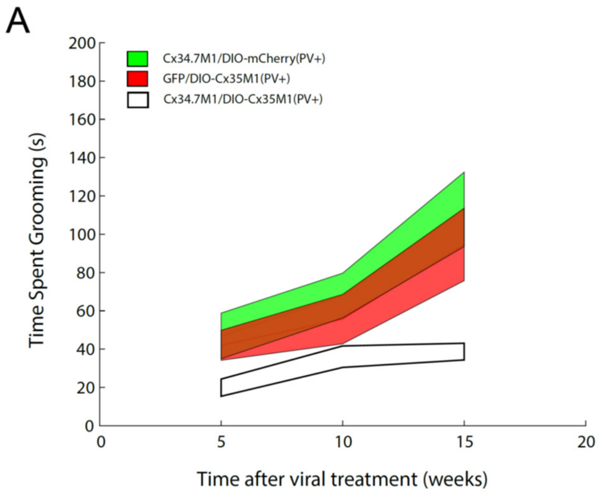


**Supplemental Figure S2: Alternative patterns of $Cx34.7_{M1}$ and $Cx35_{M1}$ expression across the OFC→Str circuit have no impact on grooming time. A)** Longitudinal quantification of grooming time in *Sapap3-/-::PV-Cre* mice with only cortical ($Cx34.7_{M1}$) or striatal PV ($Cx35_{M1}$) LinCx expression. **B)** Longitudinal quantification of grooming time in *Sapap3-/-::PV-Cre* mice with LinCx expression across all OFC→Str circuits (non-cell type specific, blue) or LinCx expression across all OFC→Str circuits except the OFC→Str PV+ circuit (purple). In both A) and B) data from mice with LinCx expression across the OFC→Str PV+ circuit are shown in white for purposes of comparison. Note that these same data are shown in Fig. 1F, and all data acquired in *Sapap3-/-::Pv-Cre* mice shown here and in Fig. 1F were analyzed together in a single statistical model; $p>0.9999$ for $Cx34.7_{M1}$/DIO-mCherry (green) and GFP/DIO-$Cx35_{M1}$ (red) at 5, 10, and 15 weeks; $p=0.0978$ at 5 weeks, $p>0.9999$ at 10 and 15 weeks for $Cx34.7_{M1}$/CamKII-$Cx35_{M1}$ (blue): $p=0.0228$ at 5 weeks, $p=0.2610$ at 10 weeks, $p>0.9999$ at 15 weeks for $Cx34.7_{M1}$/DO-$Cx35_{M1}$ (pink); all comparisons against fluorophore controls (EGFP/DIO-mCherry, see Fig. 1F) using Kruskal-Wallis multiple comparison test with Dunn's correction. Data shown as mean±s.e.m.

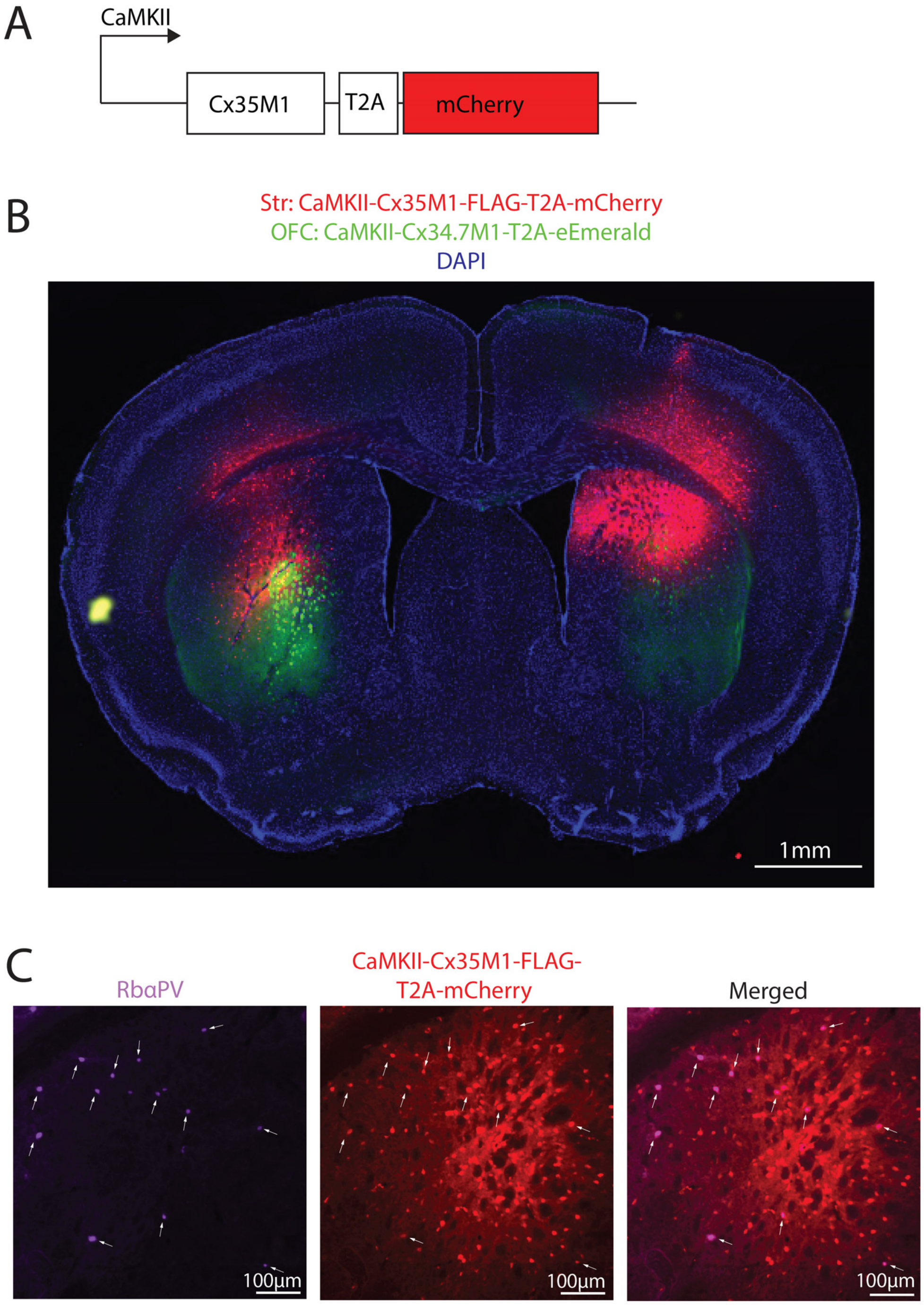


**Supplemental Figure S3: CaMKII promotor yields Cx35$_{M1}$ expression in Str PV+ interneurons and other cell types. A)** Schematic of AAV9 viral construct to target CaMKII expressing neurons. **B)** Representative coronal site showing expression of Cx35$_{M1}$-T2A-mCherry in Str (red) and Cx34.7$_{M1}$ in OFC projection neurons (green). **C)** Cx35M1-T2A-mCherry neurons colocalize with immunostaining against parvalbumin (PV). Image showing cells stained with rabbit anti-PV (purple, left), cells expressing mCherry (red, middle), and their overlap (right). White arrows indicate cells positive for both PV and mCherry.

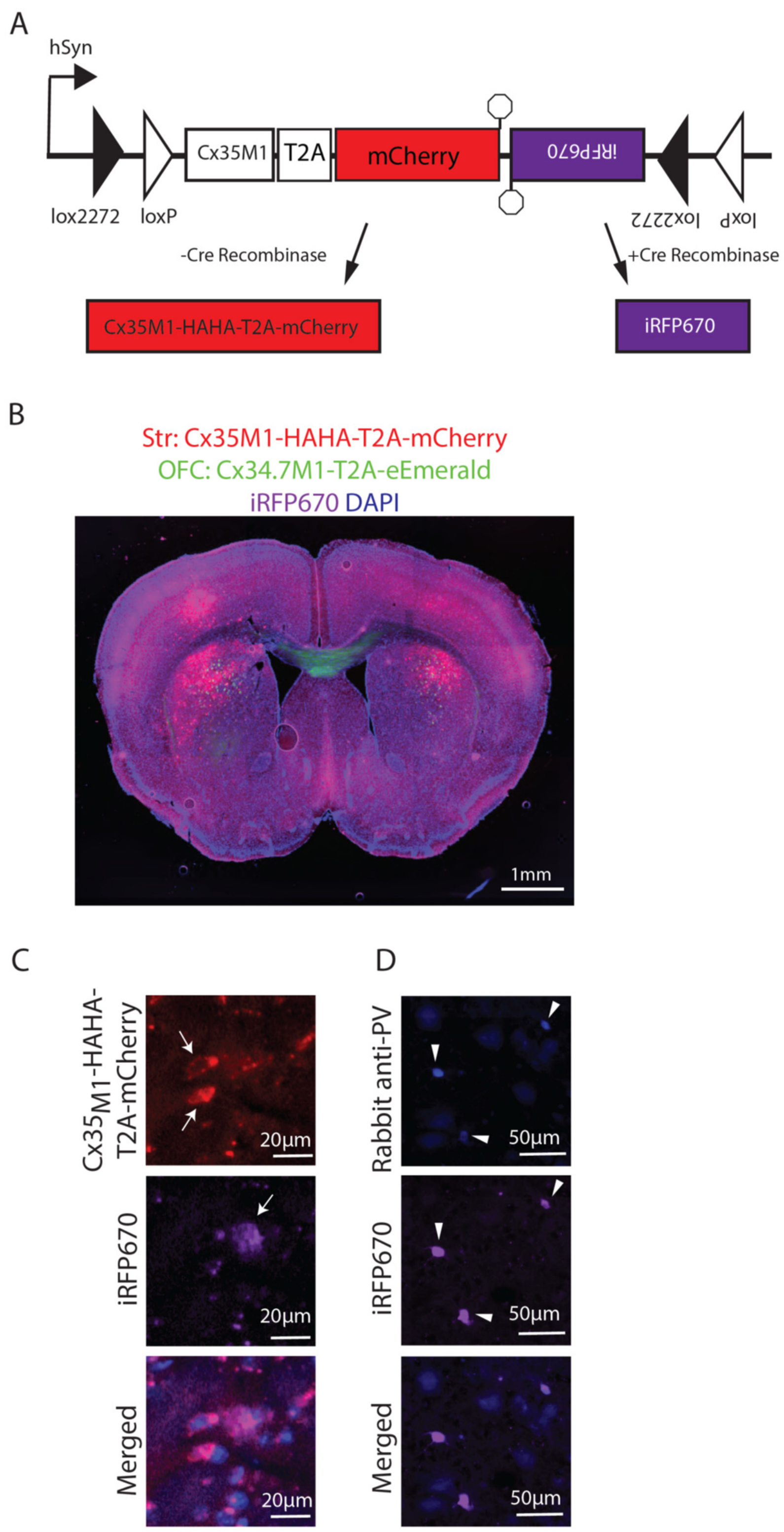


**Supplemental Figure S4: Targeting $Cx35_{M1}$ to non-PV-expressing neurons in Str.**

**A)** Schematic of AAV9 viral construct to only express $Cx35_{M1}$ in cells that do not express PV, using a Cre-switch with Cre-off $Cx35_{M1}$-T2A-mCherry and Cre-on iRFP670. **B)** Representative coronal brain slice showing expression of $Cx35_{M1}$-T2A-mCherry (red) and iRFP670 (purple) in Str and $Cx34.7_{M1}$-EGFP from OFC projection neurons (green). **C)** mCherry and iRFP670 are expressed by different neurons in *Sapap3-/-::PV-Cre* mouse injected with the DO virus. White arrows indicate neurons that express either mCherry (top) or iRFP670 (middle). **D)** iRFP670-expressing Str neurons colocalize with immunostaining against parvalbumin (Rabbit anti-PV) in *Sapap3-/-::PV-Cre* mice injected with DO virus. White arrowheads indicate positive cells for both PV and iRFP670.

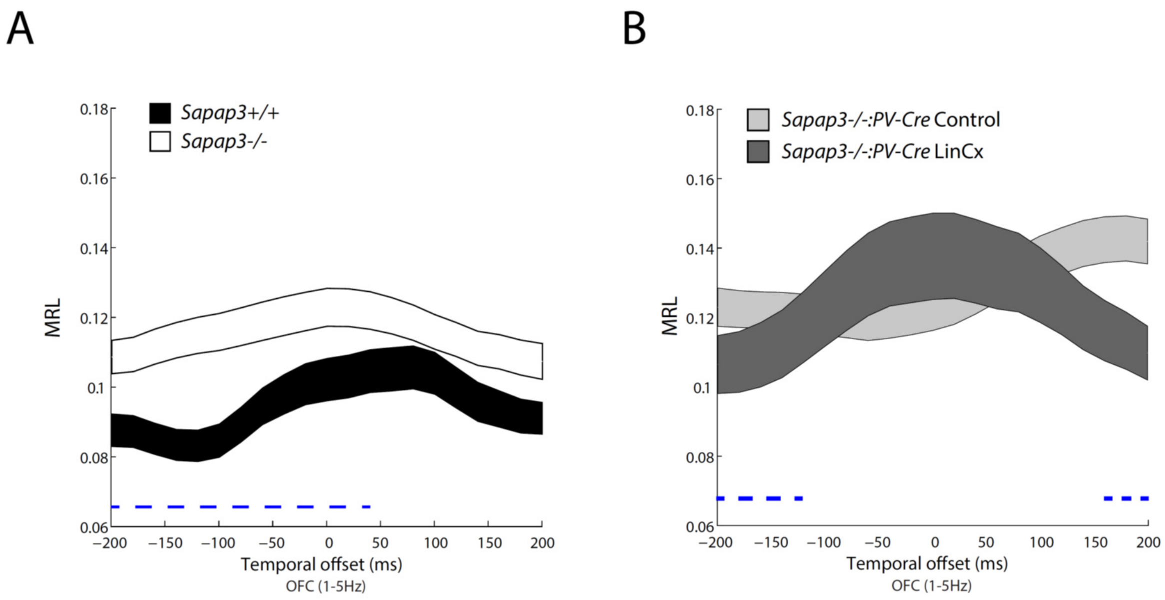


**Supplemental Figure S5: SPN timing is moderated by LinCx in *Sapap3-/-* mice. A-B)** Coupling analyses of Str SPN units to OFC oscillations at temporal offsets ranging from -200 to 200 ms. (Reanalysis of data presented in Fig. 3E-F). Blue dashed line indicates offsets which were significantly different between groups using Mann-Whitney U test with FDR correction for comparisons across the 21 offsets. Data shown as mean±s.e.m.

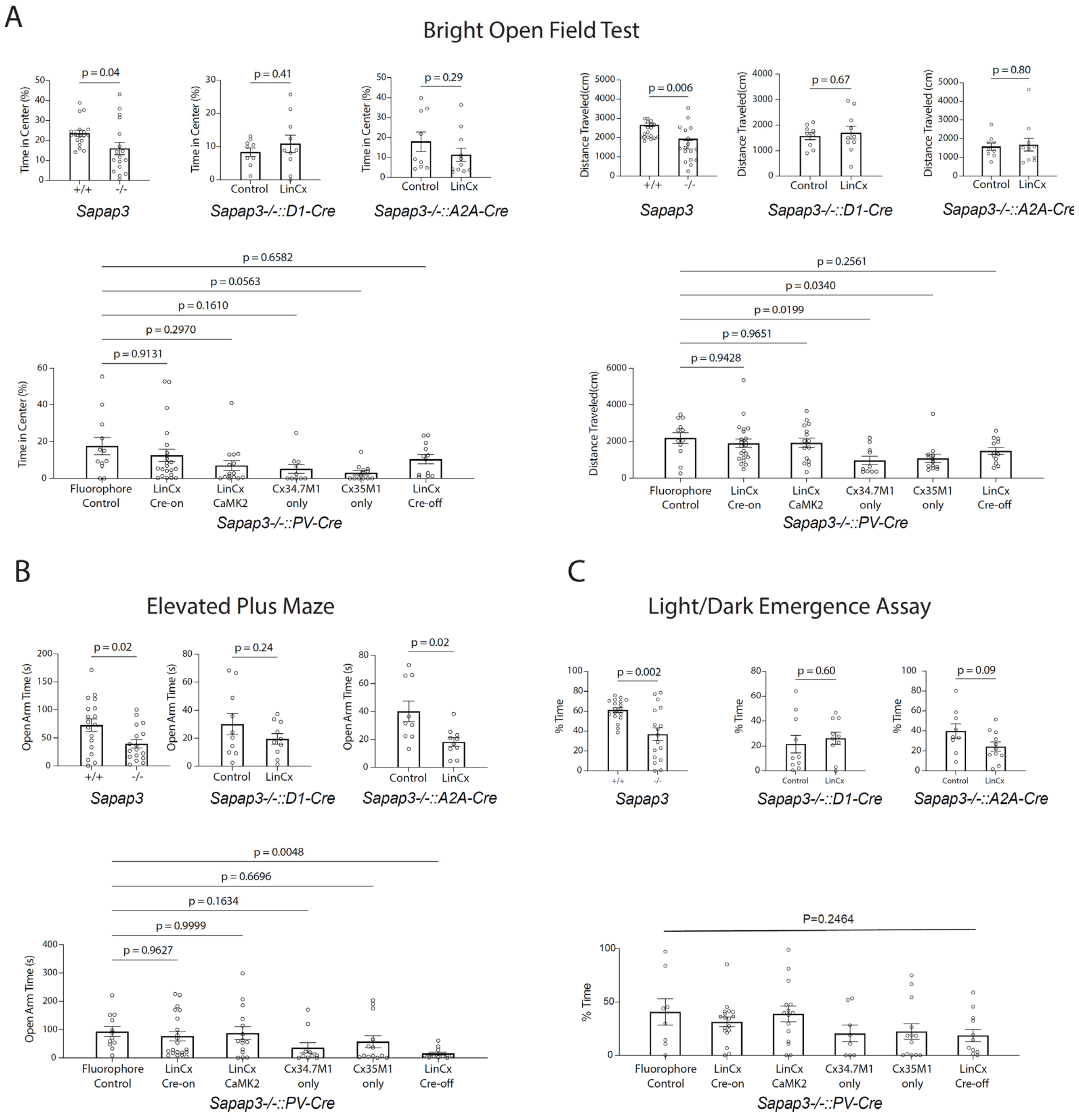


**Supplemental Figure S6: Anxiety-like behaviors in LinCx-manipulated *Sapap3-/-* mice**

**A)** (top, left 3 graphs) Percent time in center during a 5-min bright open field test in *Sapap3 +/+* and *-/-* mice (t=2.161, df=23.96, p=0.04 using an unpaired t-test with Welch's correction), D1-Cre LinCx-manipulated mice (t=1.238, df=13.41, p=0.24 using an unpaired t-test with Welch's correction), A2A-Cre LinCx-manipulated mice (t=2.722, df=10.83, p=0.02 using an unpaired t-test with Welch's correction), and (bottom left) PV-Cre LinCx-manipulated mice with Cre controls (W=3.595, p=0.0106 using a Welch's ANOVA; adjusted p-values Dunnett's T3 multiple comparison test). (top, right 3 graphs) Total distance traveled during a 5-min bright-OFT in *Sapap3 +/+* and *-/-* mice (t=3.020, df=21.66, p=0.006 using an unpaired t-test with Welch's correction), D1-Cre LinCx manipulated mice (t=0.4371, df=14.22, p=0.67 using an unpaired t-test with Welch's correction), A2A-Cre LinCx manipulated mice (t=0.2540, df=15.72, p=0.80 using an unpaired t-test with Welch's correction), and (bottom right) PV-Cre LinCx-manipulated mice with Cre controls (W=3.657, p=0.0043 using Welch's ANOVA, adjusted p-values from Dunnett's T3 multiple comparisons test included on figure). **B)** Open arm time during a 5-min elevated plus maze in *Sapap3 +/+* and *-/-* mice (t=2.468, df=29.29, p=0.02 using an unpaired t-test with Welch's correction), D1-Cre LinCx-manipulated mice (t=1.238, df=13.41, p=0.24 using an unpaired t-test with Welch's correction), A2A-Cre LinCx-manipulated mice(t=2.722, df=10.83, p=0.02 using an unpaired t-test with Welch's correction), and PV-Cre LinCx-manipulated mice with Cre controls (W=6.689, P=0.0002 using a Welch's ANOVA test, adjusted p-values from Dunnett's T3 multiple comparisons test included on figure). **C)** Percent time in light chamber during a 5-min light/dark emergence assay in *Sapap3 +/+* and *-/-* mice (t=3.500, df=20.97, p=0.002 using an unpaired t-test with Welch's correction), D1-Cre LinCx-manipulated mice (t=0.5357, df=16.11, p=0.60 using an unpaired t-test with Welch's correction), A2A-Cre LinCx-manipulated mice (t=1.810, df=14.01, p=0.09 using an unpaired t-test with Welch's correction), and PV-Cre LinCx-manipulated mice with Cre controls (W=1.428, p=0.2464 using a Welch's ANOVA test). Significant p-values based on α=0.05/3 to correct for the three assays each mouse had undergone. Data shown as mean±s.e.m.
